## Extended Data for "Lung extracellular matrix modulates KRT5^+^ basal cell activity in pulmonary fibrosis"

**Extended data Fig. 1. KRT5+ BC distribution in the normal and fibrotic distal lung.**

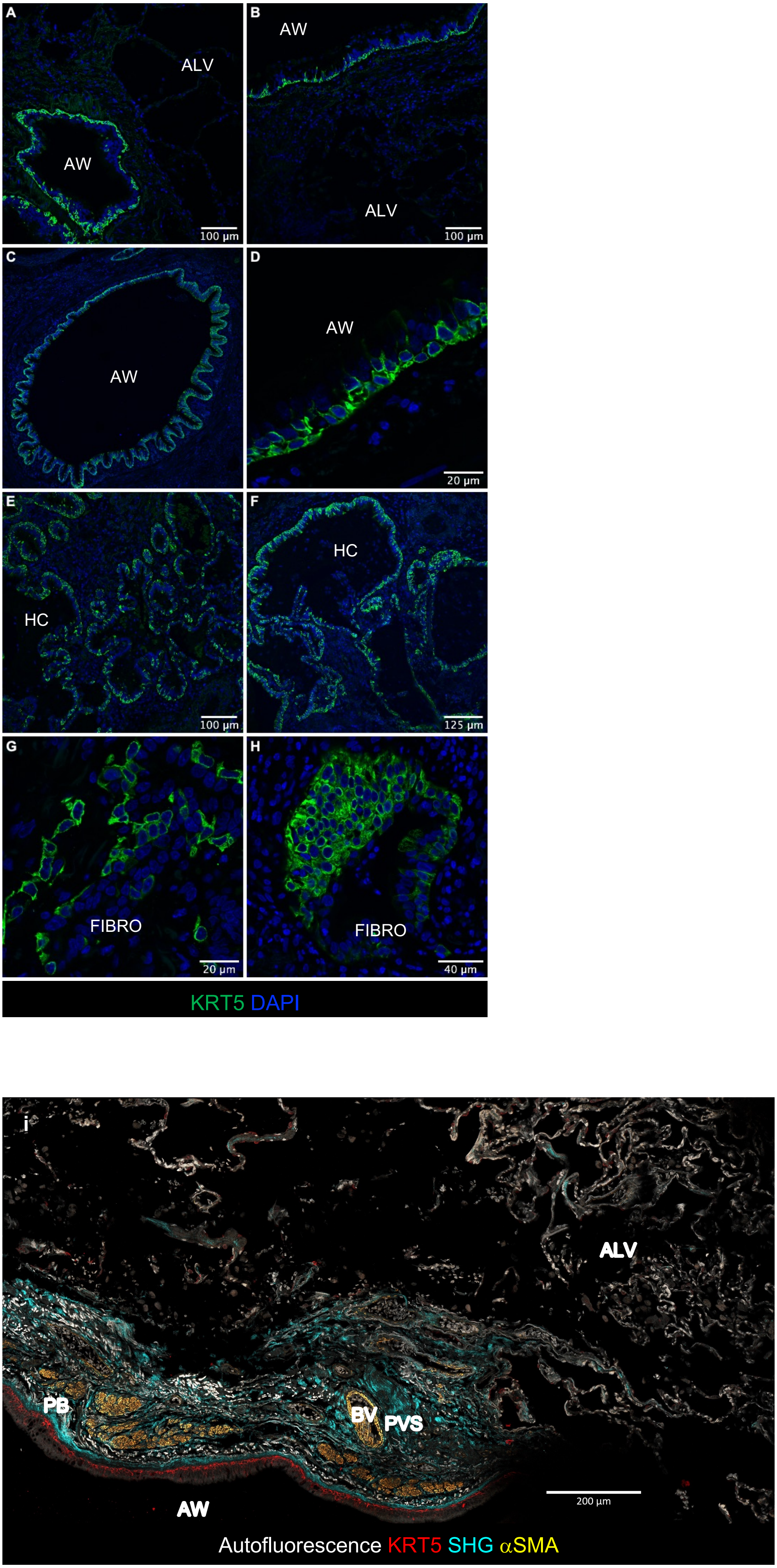

**Extended data Fig. 1. $KRT5^+$ BC distribution in the normal and fibrotic distal lung.** (a – h) Distal lung tissue sections from controls and IPF patients immunostained for basal cell marker $KRT5^+$ (green) and DAPI (blue). (a, b) In control lung tissue KRT5+ cells are found restricted to the distal airways (AW) and are absent from the alveolar tissue (ALV). (c, d) In IPF lung tissue, $KRT5^+$ cells are again seen in the distal airways in a typical configuration but are also located in (e, f) honeycomb cysts (HC), and (g, h) fibrotic interstitium (FIBRO). (i) Overview of normal distal lung tissue showing $KRT5^+$ cells (red) lining the airway but absent from the alveolar region. SHG signal (turquoise) in the peribronchial (PB) and perivascular (PVS) regions. Alpha-smooth muscle actin (yellow) outlining smooth muscle bands surrounding airways.

**Extended data Fig. 2. Quantitative image analysis of control and IPF distal lung.**

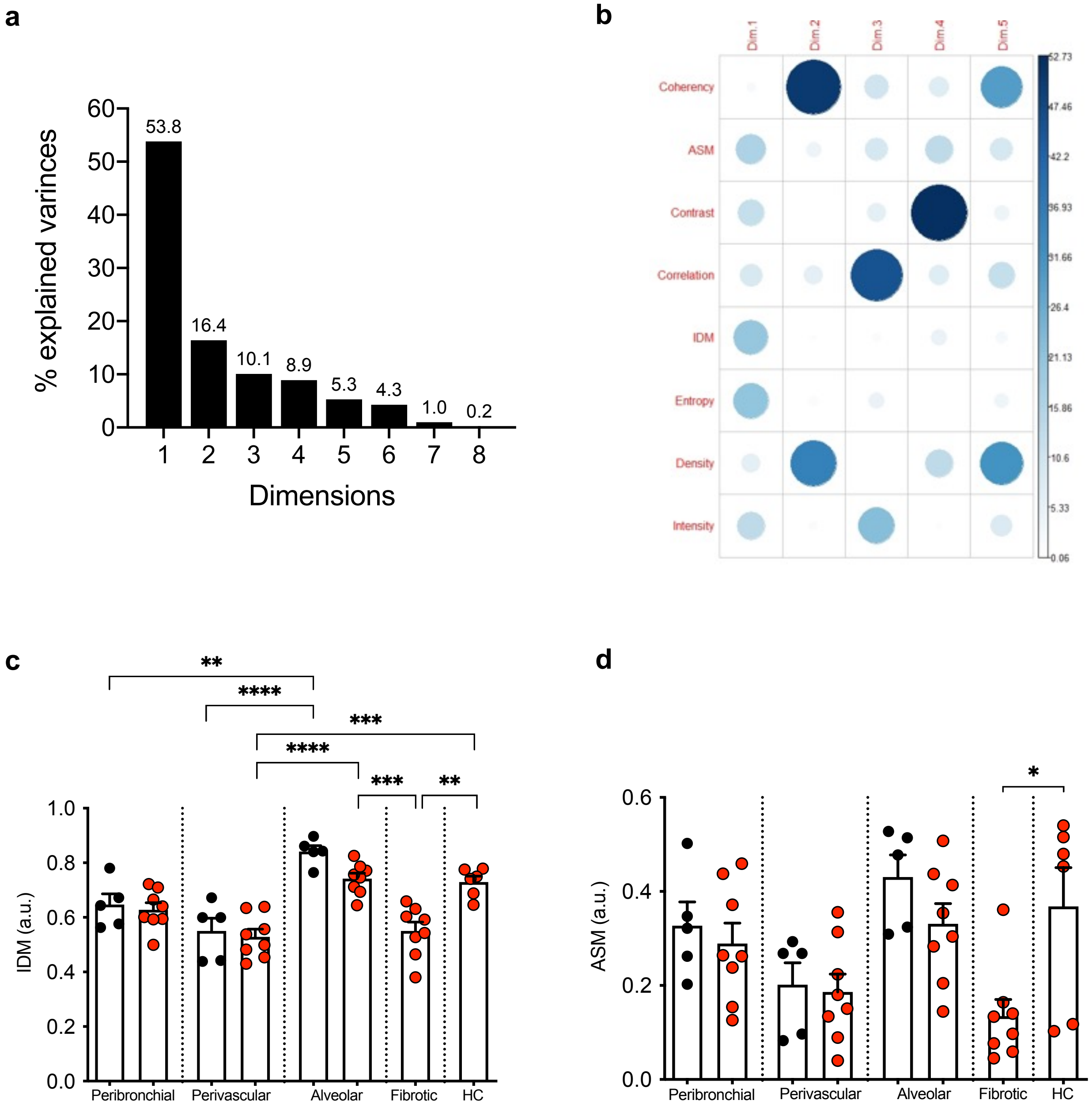

**Extended data Fig. 2. Quantitative image analysis of control and IPF distal lung** (a) Scree plot of % variances explained by each component (dimension) of PCA plot shown in Fig. 1d. (b) Corr plot of variables contributing to each dimension of PCA. (c - d) Image texture analysis per region showing (c) inverse difference moment (IDM) and (d) angular second moment (ASM). Data plotted as mean + SEM. Each data point represents the average of 2 – 6 images per area per subject. Ordinary one-way ANOVA with Tukey's multiple comparisons test ****$P$<0.0001, ***$P$<0.001, **$P$<0.01, *p<0.05.

**Extended data Fig. 3. IMC of perivascular region of distal lung tissue from controls**

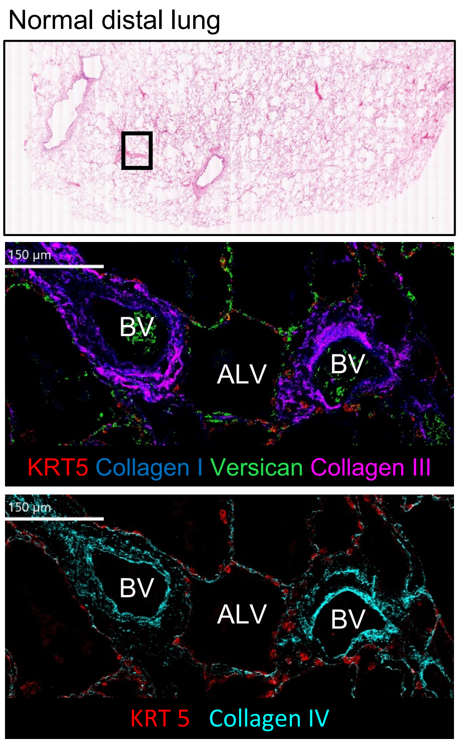

**Extended data Fig. 3. IMC of distal lung tissue from controls and patients with IPF** Imaging mass cytometry (IMC) of perivascular region of distal lung tissue from normal control (n = 1; H&E overview shown in top panel) showing distribution of ECM components; collagen I (blue), collagen III (magenta) and versican (green) (middle panel), collagen IV (turquoise) (bottom panel) in relation to $KRT5^+$ BCs (red). Representative image demonstrating key regions; blood vessel (BV) and alveolar (ALV).

**Extended data Fig. 4. KRT5+ basal cells co-localise with fibroblasts in the fibrotic niche.**

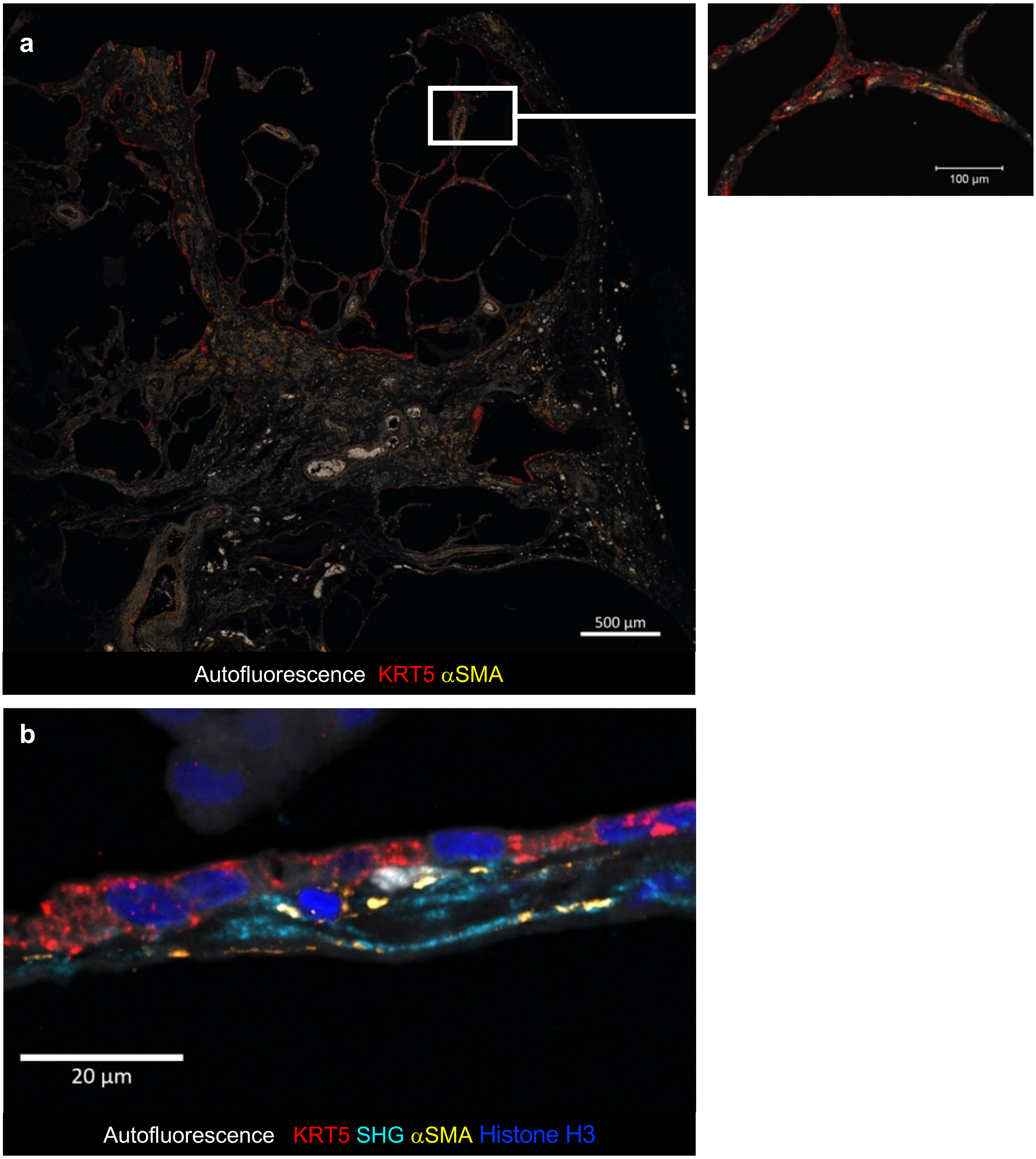

**Extended data Fig.4. KRT5+ basal cells co-localise with fibroblasts in the fibrotic niche.** (a) Overview of fibrotic, remodelled alveolar region of IPF lung with high-resolution image (white box) showing co-localisation of KRT5+ basal cells (red) with $\alpha$-SMA+ fibroblasts (yellow). (b) High resolution image of fibrotic alveolar region of IPF lung showing co-localisation of KRT+ basal cells, $\alpha$-SMA+ fibroblasts and SHG signal (cyan).

**Extended data Fig. 5. Human airway basal cell phenotyping.**

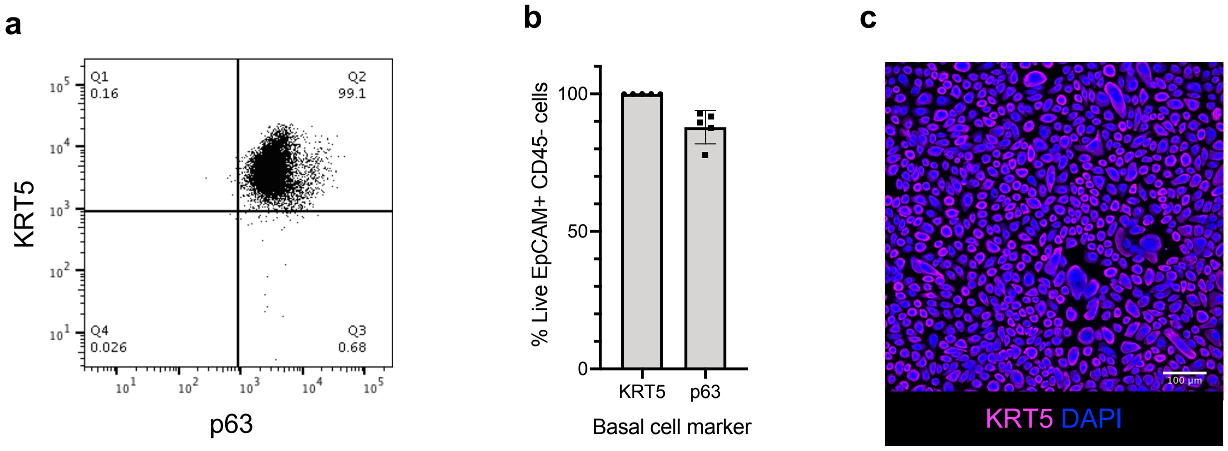

**Extended data Fig.5. Human airway basal cell phenotyping.** (a, b) Flow cytometry phenotyping of primary human airway epithelial cells in submerged culture (passage 3) for basal cell markers KRT5 and p63. (c) Immunofluorescence microscopy for KRT5 in submerged culture.

**Extended data Fig. 6. Gene expression of IPF KRT5+ cells compared to healthy KRT5+ cells cultured on collagen I or IPF CDMs.**

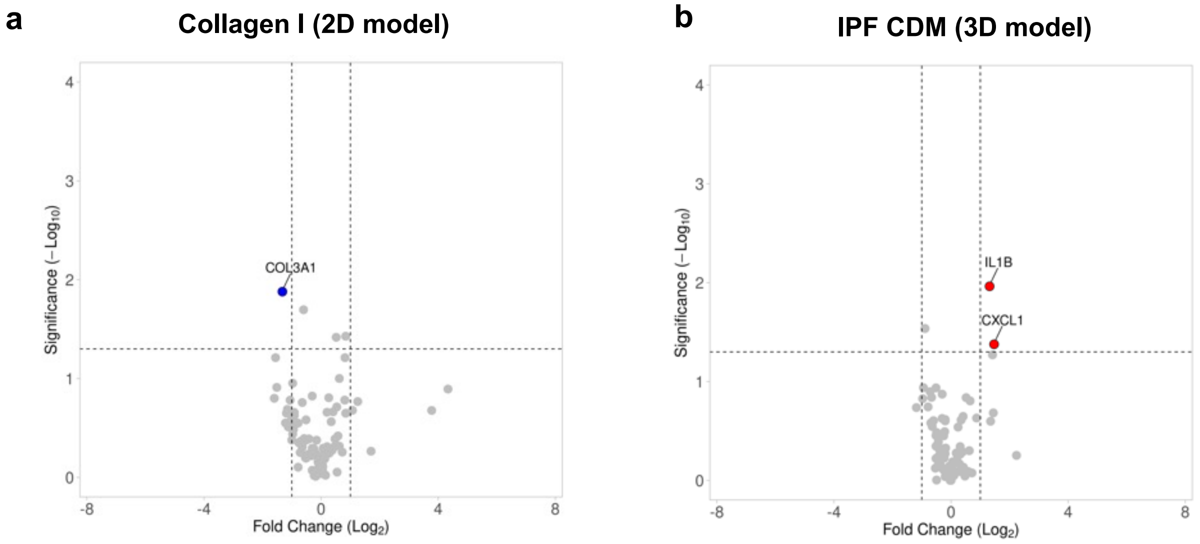

**Extended data Fig.6. Gene expression of IPF KRT5+ cells compared to healthy KRT5+ cells cultured on collagen I or IPF CDMs.** (a, b) Volcano plots showing gene expression changes determined by Qiagen $RT^2$ PCR Array between KRT5+ cells from IPF patients vs. healthy controls cultured on (a) collagen I or (b) IPF CDMs. Upregulated genes in IPF KRT5+ cells vs. control KRT5+ cells (red), downregulated genes in IPF KRT5+ cells vs. control KRT5+ cells (blue). Significance defined as log fold change >2 or <2, $p<0.05$. P values were calculated based on a Student's t-test of the replicate normalised gene expression values (2^(- Delta CT)) for each gene in control and test groups. Housekeeping genes used - ACTB, B2M and RPLP0.

**Extended data Fig.7.**

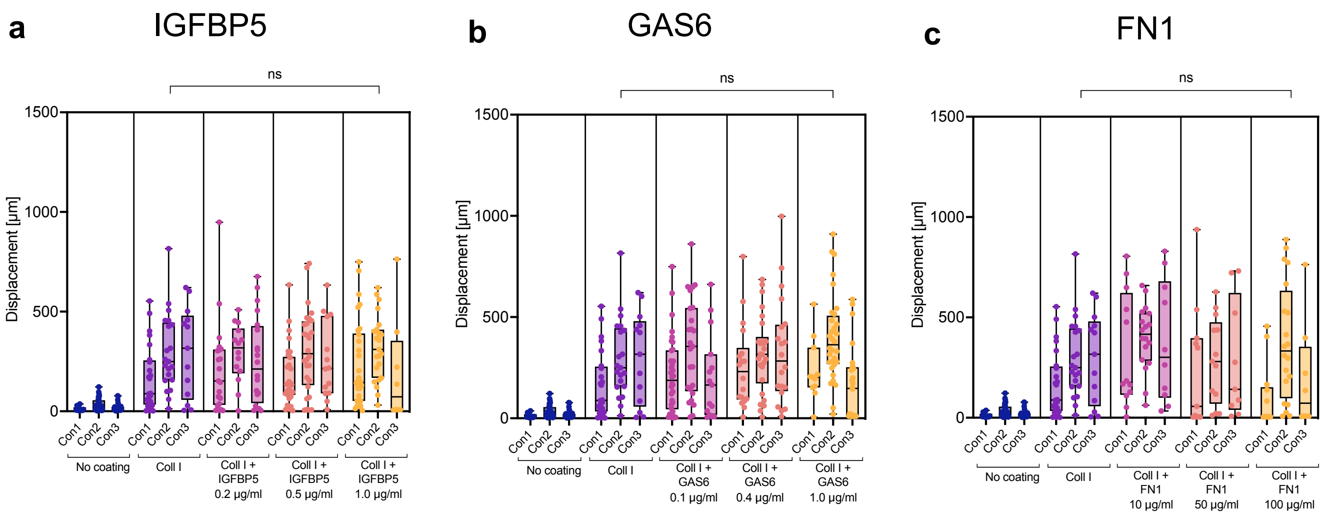

**Extended data Fig.7. Influence of proteins identified by MS-proteomics on KRT5$^+$ cell migration.** (a– c) Box and whisker dot plots quantifying displacement of cells from starting position for KRT5+ cells cultured on (a) IGFBP5; (b) GAS6; and (c) FN1. A nested one-way anova and Tukey's multiple comparison test used for statistical analysis.

**Extended data Table 1. Lung tissue sections used for image analysis experiments.**

| Sample ID | Diagnosis | Lung histology | Age (yrs) | Gender |
|---|---|---|---|---|
| CTRL_TISSUE_1 | Control | NSCLC; adenocarcinoma | 66 | F |
| CTRL_TISSUE_2 | Control | NSCLC; pleomorphic carcinoma | 74 | M |
| CTRL_TISSUE_3 | Control | NSCLC; pleomorphic carcinoma | 68 | M |
| CTRL_TISSUE_4 | Control | NSCLC; pleomorphic carcinoma | 66 | F |
| CTRL_TISSUE_5 | Control | NSCLC; adenocarcinoma | 64 | M |
| IPF_TISSUE_1 | IPF | Usual interstitial pneumonia | 64 | F |
| IPF_TISSUE_2 | IPF | Usual interstitial pneumonia | 63 | M |
| IPF_TISSUE_3 | IPF | Usual interstitial pneumonia | 64 | M |
| IPF_TISSUE_4 | IPF | Usual interstitial pneumonia | 64 | M |
| IPF_TISSUE_5 | IPF | Usual interstitial pneumonia | 59 | M |
| IPF_TISSUE_6 | IPF | Usual interstitial pneumonia | 57 | F |
| IPF_TISSUE_7 | IPF | Usual interstitial pneumonia | 69 | M |
| IPF_TISSUE_8 | IPF | Usual interstitial pneumonia | 77 | M |

**Extended data Table 2. Quantitative image analysis measurements**

| Measurement | Software | Description |
|---|---|---|
| Collagen intensity | Imaris Software, Oxford Instruments | Mean intensity of SHG signal in collagen fibres. |
| Collagen density | Imaris Software, Oxford Instruments | Collagen area represented as a percentage coverage of the total image area analysed. |
| Coherency | OrientationJ plugin, Fiji | Measures fibre orientation and coherency; a value of zero indicates perpendicular fibres, a value of 1 indicates parallel fibres. |
| Contrast | GLCM plugin, Fiji | Measures local variations in the grey-level co-occurrence matrix; a value of zero indicates a constant image with no variation. |
| Entropy | GLCM plugin, Fiji | Statistical measure of randomness, ranging from zero to infinity. |
| Correlation | GLCM plugin, Fiji | Linear dependency of grey levels on those of neighbouring pixels; higher values for similar grey-level regions. |
| Inverse Difference Moment (IDM) | GLCM plugin, Fiji | Local homogeneity or smoothness across an image; a value of 1 indicates a grey levels of the pixel pairs are similar |
| Angular Second Moment (ASM) | GLCM plugin, Fiji | Measures the number of repeated pixel pairs indicating uniformity of distribution of grey level in the image; a value of 1 indicates a constant image. |

**Extended data Table 3. Catalogue of human primary cells used for the study.**

| | Sample | Cell type | Diagnosis | Age (yrs) | Gender | Smoking status |
|---|---|---|---|---|---|---|
| **Fig. 3** | Con1 | KRT5+ BC | Healthy | 67 | M | NA |
| | Con2 | KRT5+ BC | Healthy | 65 | M | NA |
| | Con3 | KRT5+ BC | Healthy | 63 | M | Ex |
| | Con4 | KRT5+ BC | Healthy | 55 | F | Never |
| | Con5 | KRT5+ BC | Healthy | 63 | M | Never |
| | IPF1 | KRT5+ BC | IPF | 53 | M | Current |
| | IPF2 | KRT5+ BC | IPF | 67 | M | Current |
| | IPF3 | KRT5+ BC | IPF | 78 | M | Ex |
| | IPF4 | KRT5+ BC | IPF | 76 | F | Ex |
| | IPF5 | KRT5+ BC | IPF | 68 | M | Ex |
| **Fig. 4** | Con1 | KRT5+ BC | Healthy | 26 | F | Ex |
| | Con2 | KRT5+ BC | Healthy | 51 | M | Ex |
| | Con3 | KRT5+ BC | Healthy | 50 | F | Never |
| | Con4 | KRT5+ BC | Healthy | 47 | M | Ex |
| | IPF1 | KRT5+ BC | IPF | 72 | M | Ex |
| | IPF2 | KRT5+ BC | IPF | 73 | M | Never |
| | IPF3 | KRT5+ BC | IPF | 80 | M | Never |
| | IPF4 | KRT5+ BC | IPF | 80 | F | Never |
| | IPF HLF for CDM | HLF | IPF | 67 | F | NA |
| **Fig. 5** | Con1 | KRT5+ BC | Healthy | 67 | M | Ex |
| | Con2 | KRT5+ BC | Healthy | 47 | M | Ex |
| | Con3 | KRT5+ BC | Healthy | 26 | F | Ex |
| | IPF1 | KRT5+ BC | IPF | 52 | M | Ex |
| | IPF2 | KRT5+ BC | IPF | 80 | M | Never |
| | IPF3 | KRT5+ BC | IPF | 72 | M | Ex |
| | IPF HLF for CDM | HLF | IPF | 67 | F | NA |
| **Fig. 6** | Con1 | HLF | Normal tissue, carcinoid | 65 | F | Never |
| | Con2 | HLF | Normal tissue, adenocarcinoma | 64 | M | Never |
| | Con3 | HLF | Normal tissue, adenocarcinoma | 66 | M | Ex |
| | IPF1 | HLF | IPF | 67 | F | NA |
| | IPF2 | HLF | IPF | 51 | M | Never |
| | IPF3 | HLF | IPF | 62 | F | NA |
| | Healthy KRT5+ BC | KRT5+ BC | Healthy | 67 | M | Ex |
| **Fig. 7** | Con1 | KRT5+ BC | Healthy | 50 | F | Never |
| | Con2 | KRT5+ BC | Healthy | 47 | F | Never |
| | Con3 | KRT5+ BC | Healthy | n/a | n/a | n/a |
| | Con4 | KRT5+ BC | Healthy | 63 | M | Ex |
| | Con5 | KRT5+ BC | Healthy | 55 | F | Never |
| | Con6 | KRT5+ BC | Healthy | 63 | M | Never |

**Extended data Video 1.** KRT5$^+$ BC migration varied according to ECM ligand.

**Extended data Videos 2 - 4.** KRT5$^+$ BCs from healthy controls and IPF patients were fluorescently labelled and tracked on the fibrotic CDMs.
